# Aberrant accumulation of α-synuclein might be linked with the progressive motor deficits in a mouse model of Angelman syndrome

**DOI:** 10.64898/2026.08.27.747679

**Authors:** Sagarika Das, Bhaskarjyoti Giri, Nihar Ranjan Jana

**Affiliations:** Neurobiology of Disease Laboratory, Department of Bioscience and Biotechnology, Indian Institute of Technology, Kharagpur-721302, India

**Author notes:** Address correspondence to: Nihar Ranjan Jana, Department of Bioscience and Biotechnology, Indian Institute of Technology, Kharagpur-721302, India.

**Keywords:** Ube3a, Angelman syndrome, α-synuclein, Parkinson’s disease, Striatum

## Abstract

Dysfunction of maternal UBE3A leads to Angelman syndrome (AS), which is characterized by significant intellectual and motor debilities. However, the molecular underpinnings of the behavioral deficits associated with UBE3A dysfunction remain obscure. In this study, we utilized a model mouse of AS and report, for the first time, that the aberrant accumulation of α-synuclein may be linked to the development of AS. Firstly, we demonstrated a progressive deterioration of various motor functions in AS mice beginning from the early adolescent phase. Subsequently, we observed an age-dependent increase in the accumulation of both soluble and insoluble α-synuclein, including its pathological variant (pSer129), in the striatum and substantia nigra dopaminergic neurons of AS mice. We also found that Ube3a interacts with α-synuclein and promotes its proteasome-mediated degradation, as evidenced by decreased levels of K48-linked polyubiquitinated α-synuclein in the brain samples of AS mice in comparison to wild-type animals. Finally, using an RT^2^ Profiler PCR Array that analysed 84 genes specifically related to dopamine and serotonin pathways, we identified altered transcript level of various genes in the striatal tissues of AS mice that are commonly associated with nigrostriatal dysfunctions in Parkinson’s disease. These findings highlight α-synuclein as a novel target of Ube3a and suggest that α-synuclein pathology may contribute to the progressive motor and other behavioral abnormalities witnessed in AS mice.

## Introduction

Angelman syndrome (AS) is a neurodevelopmental disorder clinically represented by substantial intellectual debilities, speech deficiencies, motor deficits, periodic seizures, and sleep disturbances. Persons affected with AS also display distinctive behavioral attributes, like frequent laughter, hand-flapping gestures, hyperactivity, and a profound attraction to water [1–4]. The primary cause of AS is the deletion of the maternal chromosome 15q11-q13 locus, along with other genetic abnormalities, which ultimately results in the lack of expression of the maternally inherited *UBE3A* gene at this chromosomal position [5,6,3]. This locus of the chromosome is distinctively controlled by genomic imprinting in a tissue-selective and parent-of-origin fashion. Notably, the expression of paternal *UBE3A* gene is epigenetically repressed in the mature neuron with the generation of a relatively long noncoding antisense RNA copy, UBE3A-ATS, even though it is biallelically transcribed in other cell types[7,8]. As a result, neurons primarily rely on maternal-specific expression of the *UBE3A* gene to function properly. Likewise, the murine brain exhibits paternal-specific imprinting of the *Ube3a* gene, and the AS mouse model generated through disruption of the maternal *Ube3a* allele reproduces numerous behavioral insufficiencies observed in AS, indicating a crucial role for Ube3a in disease development [9–13].

The *Ube3a* gene transcribes and translates a 100 kDa protein and was first functionally categorized as an E3 ubiquitin ligase, an enzyme in the ubiquitination cascade that selectively recognizes substrates for polyubiquitination and subsequent elimination via either the proteasome or autophagy[14]. Consequently, Ube3a is also characterized as a transcriptional coactivator of several nuclear hormone receptors[15]. Therefore, it is presumed that AS phenotypes are triggered by the absence of both ubiquitin ligase and coactivator activity of Ube3a. To date, a number of Ube3a substrates have been identified, and many of them are essential in regulating synaptic maturation, function, and plasticity [16–20]. Unusual accumulations of various substrates in different brain regions of AS mice are implicated as underlying causes of synaptic dysfunction, destabilization of the neuronal network, and resulting behavioral deficits[21,20,18]. On the other hand, the importance of Ube3a’s coactivator function in AS pathogenesis is poorly understood, despite Ube3a being predominantly a nuclear protein, and despite the evidence that absence of nuclear Ube3a could be the principal cause of disease pathogenesis[22,23].

The motor dysfunctions displayed by AS patients, like tremors, bradykinesia, rigidity, etc., closely resemble Parkinson’s disease (PD). In fact, a case study of two adults with AS showed that levodopa treatment improved motor abnormalities[24]. Subsequent studies in the AS mouse model demonstrated alterations in dopaminergic signaling and reduced function of dopaminergic neurons in the nigrostriatal pathway[25–27]. Surprisingly, these AS mice also exhibit increased dopamine release in the mesolimbic pathway [26]. Ube3a has also been shown to modulate dopamine synthesis by regulating GTP cyclohydrolase-1, possibly through its coactivator activity [28]. However, a double-blind randomized clinical trial in AS children with levodopa failed to improve motor deficits, which prompted us to investigate further impaired dopaminergic function in the brains of AS mice[27]. Interestingly, Ube3a was earlier shown to promote the degradation of wild-type α-synuclein and some of its PD-associated mutants[29]. However, the mechanism by which Ube3a regulates α-synuclein clearance and, most importantly, the role of α-synuclein in the development of motor deficits in the AS mouse model remain unknown. In the present investigation, we demonstrate that age-dependent accumulation of α-synuclein and its pathological variant (pSer129) in the substantia nigra dopaminergic neurons and striatum of AS mice is linked to their dysfunction and progressive motor deterioration.

## Materials and Methods

### Materials

Mouse monoclonal antibodies raised against α-synuclein (Cat# sc-69977), Ube3a (Cat# sc-166689), Darpp32 (Cat# sc-271111), and GAPDH (Cat# sc-32233) were procured from Santa Cruz Biotechnology. Rabbit anti-α-synuclein antibody (Cat# PA1-18264) was purchased from Invitrogen. Rabbit antibodies against phosphorylated α-synuclein at Ser129 (pS129; Cat# ab51253), as well as rabbit monoclonal antibodies specific for K48-linked ubiquitin chains (EP8589; Cat# AB140601) and K63-linked ubiquitin chains (EPR8590-488; Cat# AB179434), were purchased from Abcam. Rabbit polyclonal antibody against tyrosine hydroxylase (Cat# AB152), the reverse transcription (RT) kit (Cat# 4368814), and SYBR Green Master Mix (Cat# A25742) for quantitative PCR were acquired from Sigma-Aldrich. Immobilon Western Chemiluminescent Horseradish Peroxidase (HRP) Substrate (Cat# WBKLS0500) was obtained from Merck. Fluorescent-tagged secondary antibodies were sourced from Thermo Fisher Scientific. The RT² First Strand Kit (Cat# 330401) and RT² Profiler PCR Array for mouse dopamine and serotonin pathway-specific genes (Cat# 330231) were procured from Qiagen.

### Animals

Maternal deficient Ube3a*m-/p+* (RRID: IMSR_JAX:016590) was obtained from The Jackson Laboratory and kept in standard IVC cages with a sufficient quantity of pelleted food and water, and maintained in the animal facility of the Indian Institute of Technology, Kharagpur. Ube3a*m-/p+* females were paired with wild-type males, and the subsequent pups produced were genotyped by extracting genomic DNA from the tail. Both male and female mice were used in most our experiments. All experimentations were permitted by the Institutional Animal Ethics Committee (IAEC) of IIT Kharagpur (protocol numbers: IE-3/NJ-BS/2.22).

### Animal behavioral studies

Behavioral experiments were conducted in a blinded manner between 9 am and 5 pm, roughly at the same time each day. All behavioral tests were recorded through a video camera system and later analyzed. Every apparatus was wiped with 10% ethanol between subjects during experiments to prevent any internal olfactory cues.

#### Gait analysis

For footmark gait investigation, mice were allowed to walk inside a glass pathway (40 cm x 5 cm) placed on white paper. The hind paws of each mouse were painted red using a nontoxic paint for the purpose of analyzing the gait patterns. After two days of training, final experiments were conducted, and each mouse underwent two testing sessions. The hind limb footprint of each animal on the paper was analyzed for stride length and width. *Clasping behavior test*-The animals were suspended through gently holding their tails in the air around 10-20 cm above the animal cage for 30 sec. The time to show the clasping behavior of the hind limbs for each mouse was noted. Clasping scores were designated as follows: score 1, 15-20sec; score 2, 10-15sec; score 3, 5-10sec; and score 4, 0-5sec.

#### Pole test

For the pole test, a 50 cm-long, 1.5 cm-diameter wooden rod was used. Mice were first acclimatized by placing the animal’s head upward on the pole top and allowing them to descend. Mice were then trained for two consecutive days with two test trials on each day. In the actual test, each animal performed 3 trials; the time to incline for every single trial was noted, and the average of the 3 tests was taken for final data analysis. Finally, the time to descend by wild-type and AS mice was compared.

#### Beam balance test

A beam of 80 cm in length and 20 mm in width that was positioned 50 cm above the ground level was used to train mice. A black box was introduced at one end of the beam. As the animal moved over the beam, the number of hind-paws slides and the traversal time were recorded. Mice were first trained for two days, first with square and then circular beams (8 and 12 mm in diameter) after being taught to walk on a square beam having diameter of 20 mm. On testing day, each mouse undergoes 3 trials for each beam type at different diameters. On the first testing day, all square beams were used; on the second, all round beams. Mean values from all three trial sessions were used for the analysis.

#### Rotarod test

Mice were first trained to maintain a steady pace of 5 rpm by placing them on a rotarod until they could remain for a minimum of 30 seconds; those that fell were continually returned back on the rotarod till they were competent to stay on the rotating rod for around 30 seconds. The animals were taught for two days prior to the test, and on the test day, they were given three trials, up to a maximum speed of 25 rpm. The time taken to drop was documented, and the average across the three trials was reported. Maximum trial time was set at 150s.

### Immunofluorescence study

The animals were sedated by introducing Xylazine (10 mg/kg body weight) and Ketamine (100 mg/kg body weight), and transcardially perfused with PBS followed by 4% paraformaldehyde (PFA) in PBS. The brains were then carefully harvested and preserved in 4% PFA at 4°C. The brain samples were serially dipped in 10%, 20%, and 30% sucrose solutions for 24 hours at 4°C, then subjected to coronal sectioning (20 μm thick) using a cryotome. The serial sections were preserved in PBS comprising 0.02% sodium azide at 4°C. For immunofluorescence study, brain sections from the midbrain regions (Bregma −2.50 to −3.88 mm) were first exposed to antigen recovery solution (10 mM citrate buffer; pH 6.0), then cooled in the same retrieval solution for approximately 20 minutes. The sections were then permeabilized with 0.3% Triton X-100, blocked with 5% BSA comprising 0.1% Triton X-100, and incubated for about 12-15 hours at 4°C with primary antibodies against tyrosine hydroxylase (TH, 1:3000 dilution), α-synuclein (1:50 dilution), and Darpp32 (1:200 dilution). The sections were washed with PBS and incubated with suitable secondary antibody (1:500 dilutions) at room temperature for about 1 hour. Finally, the sections were washed several times with PBS, mounted on coverslips, and pictures were obtained using a confocal microscope (Olympus).

### Co-immunoprecipitation, immunoblotting, and dot-blot experiments

Animals were sacrificed by cervical displacement, various brain areas (cortical, hippocampal, and striatal regions) were dissected and immediately placed into liquid N_2_, and then ppreserved at −80°C until further use. Equal quantities of tissue were lysed on ice in RIPA buffer containing 50 mM Tris-HCl (pH 7.4), 150 mM NaCl, 0.1% SDS, 1% Triton X-100, 0.5% sodium deoxycholate, 0.1 mM sodium orthovanadate, 10 mM sodium fluoride, and complete protease inhibitor cocktail. The lysed samples were centrifuged at 14,000 × g for 15 minutes, soluble fractions were obtained, and protein concentrations were determined in all samples using the BCA method. The samples were then kept at −80°C in several stocks. An equal quantity of each striatal sample was mixed with SDS-PAGE sample buffer, boiled for 5 minutes, and then separated by SDS-PAGE. For co-immunoprecipitation experiments, pre-cleared cortical lysates (with protein-G agarose beads) were incubated 12-15 hours at 4°C with the primary antibodies (either Ube3a or α-synuclein), and finally incubated with protein-G agarose beads for 7–8 hours. Immunoprecipitated complexes were then washed several times with RIPA buffer, boiled with sample buffer (1X), and subjected to SDS-PAGE. The resolved proteins in SDS-PAGE were then transported onto a nitrocellulose membrane, assessed by Ponceau staining, and blocked with 5% skimmed milk for 1 hour at room temperature. The membranes were then incubated overnight (approximately 12 hours) at 4°C with primary antibodies. The following dilutions of primary antibodies were used: Ube3a, α-synuclein, and GAPDH at 1:5000; Darpp32 at 1:2000; TH at 1:10,000; K48-ubiquitin and K63-ubiquitin at 1:1,000. After several time washings with TBST, the membranes were incubated for 1 hour at room temperature with HRP-conjugated appropriate secondary antibodies (1:5000 dilutions). The immunoreactive bands were visualized using ECL reagents.

For dot-blot experiments, cortical tissues from wild-type and AS mouse were homogenized in RIPA buffer at 4°C, sonicated under ice, and then quantified by BCA. An identical amount of each sample was mixed with DNase I for 10 minutes and subsequently spun at 15,000 × g for 15 minutes. The precipitated fractions were dissolved in 2% SDS for 5 minutes under boiling condition. An equal volume of various samples (having equal amounts of protein) was passed through a nitrocellulose membrane in a dot-blot apparatus. Membranes were subsequently washed with SDS wash buffer, and preceded for immunoblotting using α-synuclein and GAPDH antibodies.

### RT² Profiler PCR Array

Total RNA was extracted from the striatal samples with the use of TRIzol reagent as per the manufacturer’s instruction, quantified, and assessed for RNA integrity through agarose gel electrophoresis. Equal amounts of total RNA (500 ng) from different samples were reverse transcribed into cDNA, diluted appropriately and added to RT² Profiler PCR Array plates along with RT² SYBR green qPCR master mix as per the supplied protocol. The cycle threshold (Ct) value from the PCR reaction were then transported to an Excel file, and the values were assessed in the web-based GeneGlobe data analysis center (https://geneglobe.qiagen.com/us/analyze) using the 2^-ΔΔCt method. In short, after uploading the Ct value, the software at initially evaluates sample quality check (PCR array reproducibility, reverse transcription efficacy, and genomic DNA contamination). After that, the Ct cut-off value (default: 35) was applied and normalized to the GAPDH, and then the system spontaneously calculates 2^-ΔΔCt and fold variation.

### RT-qPCR analysis

Extracted total RNA from the striatal tissues were subjected to cDNA synthesis using the RT kit. The qPCR was performed using SYBR green master mix on the Applied Biosystems QuantStudioTM 6 Flex Real-Time PCR apparatus. The PCR conditions were an initial denaturation at 95°C for 10 minutes, afterwards 40 cycles of denaturation at 95°C for 15 seconds, annealing at 60°C for 1 minute, and extension at 60°C for 1 minute. GAPDH was employed as a reference to standardize the mRNA levels. Fold alteration of various mRNAs were calculated with the help of the 2^-ΔΔCt technique (with the use of QuantStudio Design and Analysis Software v2.x). Primer sequences used to amplify various mRNAs are given in Supplementary Table S1.

### Statistical Analysis

All investigational data were assessed utilising GraphPad Prism software (version 8). Two-way analysis of variance (ANOVA), afterward a Bonferroni *post hoc* test, was conducted to evaluate the behavioral data. Mixed-effects analysis, afterwards a Bonferroni *post hoc* test, was used to evaluate all immunoblot data. The unpaired two-tailed Student’s *t*-test was used to calculate gene expression data. In all experimentations, values were shown as mean ± SEM, and *P*<0.05 was regarded statistically significant.

## Results

### Progressive deterioration of motor functions in AS mice

The model mouse for AS used in this study exhibits motor deficits, which seem to arise from region-specific alterations in both cerebellar and nigrostriatal dopaminergic pathways [25,30]. Using a series of behavioral tasks, we previously demonstrated that motor abnormalities in 7-8-month-old AS mice could be partly due to dysfunction of the nigrostriatal dopaminergic system[25]. To explore motor deficits in the early developmental phase and their progressive nature, we subjected AS mice, along with wild-type controls, to a series of motor tests. The tests were initiated at 1 month of age and continued until 12 months of age. We first assessed general motor function and coordination through rotarod performance and gait analysis. In the rotarod test, AS mice performed poorly compared to wild-type counterpart at 1 month age and deteriorate progressively over time (Fig. 1A). Gait investigation also showed significantly increased hind-paws stride length in AS mice matched to control group at 1 month age and gradually increased as they are getting older (Fig. 1B). A 12 months old AS mice displayed worsen rotarod performance and abnormal gait when compared to their 1 month age data. Mice were further subjected to clasping, pole, and balance beam tests to evaluate possible aberrations in nigrostriatal function. In the clasping test, AS mice took a shorter duration of hind-limb grasping (increased clasping score) at 1 month of age, which increased gradually with age (Fig. 1C). Wild-type animals did not exhibit such clasping behavior even at their age of 12 months. Comparable results were witnessed in the pole test, where AS mice took longer time to incline the vertical pole than control animals at 1 month of age. The descent time increased steadily with age and consistently showed significant differences relative to the control group (Fig. 1D). In the balance beam test, fine motor skills were compared between AS and wild-type animals. It was observed that the time consumed by the AS mice to cross both the round and square beams progressively increased with age, matching that of the wild-type group (Fig. 1E and F). The number of hind-limb slips in both square and round beam tests was also significantly more in the case of AS mice at 1 month of age. The number of hind-limb slips also progressively increased with age in AS mice (Fig. 1G and H). Results from all these motor tests indicate that motor deficits in AS mice are evident at least 1 month after birth and progressively worsen with age.

**Fig. 1.**
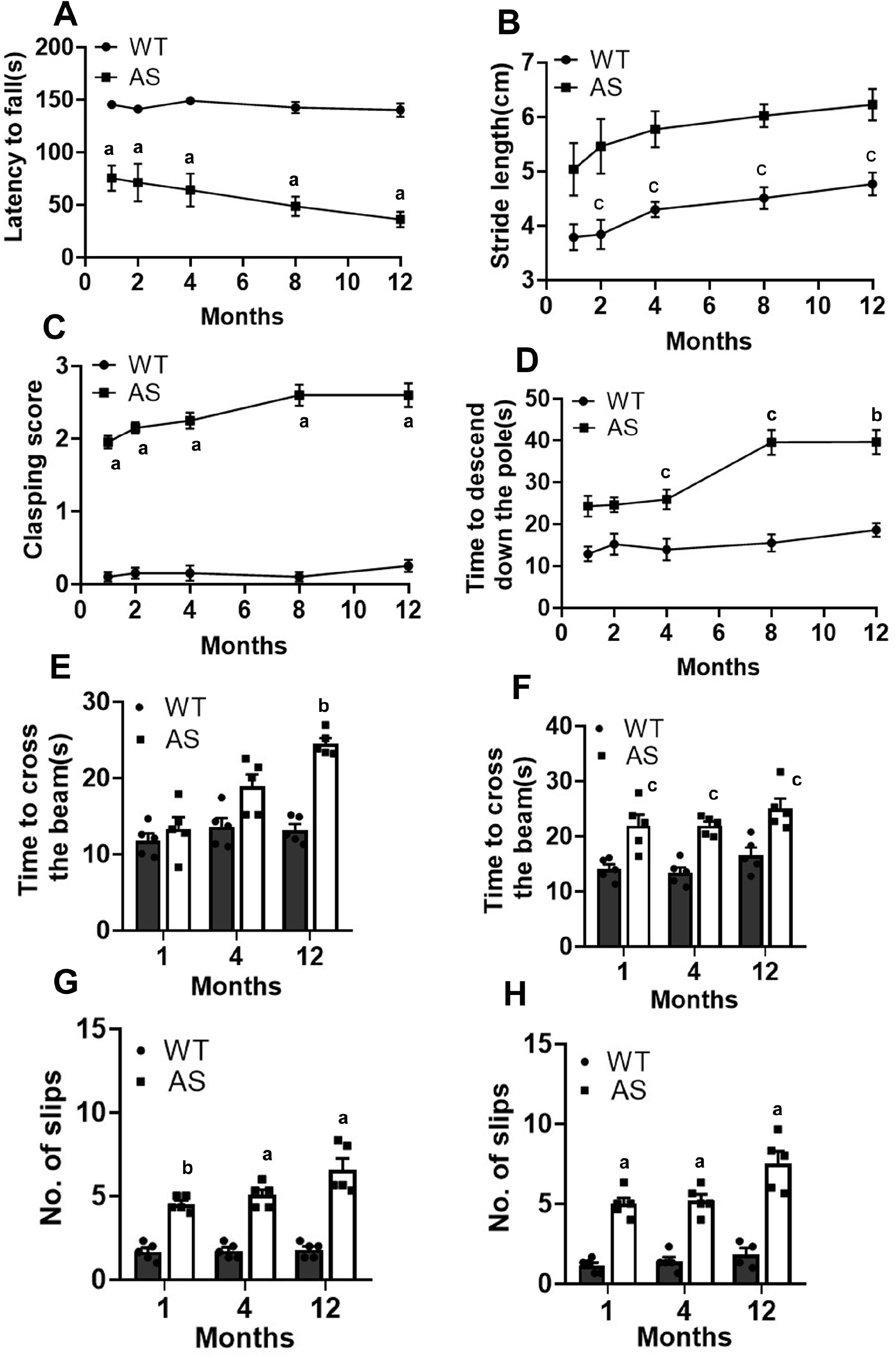
AS mice manifested progressive motor deficits. The AS mice, together with age-matched WT animals, were subjected to various motor tests starting at 1 month and repeated at different time points up to 12 months, as indicated in the figure. Experiments were initiated at the beginning of the test month and completed within a week. All of these tests recorded a slow progressive worsening of various motor functions in AS mice. A) Rotarod performance test showing latency to fall. B) Gait analysis presenting hind-limb stride length. C) Clasping behavior indicating the time taken to clasp the hind limbs. D) Pole test demonstrating the time to descend from the top of the pole. E, F) Time taken in square (E) and round (F) beams. G, H) Number of hind-limb slips in square (G) and round (H) beams. Values depicted are mean ± SEM with five animals in every investigational set. Two-way ANOVA with a Bonferroni post hoc check was used to scrutinize the data across all experiments. In the figure, a, b, and c represent *P*<0.001, *P*<0.01, and *P*<0.05, respectively, matched with the relevant WT age set. In the rotarod (*P*<0.05), clasping (*P*<0.01), and pole tests (*P*<0.05), 12-month-old AS mice exhibited significant deterioration compared to their 1-month-old counterparts. In the beam balance test, the time taken to cross the square/round beam (*P*<0.05) and the number of slips on both beams (*P*<0.01) in 12-month-old AS mice were significantly higher than at 1 month of age.

### The AS mouse brain exhibits an aberrant increase in α-synuclein and its pS129 variant levels

Since AS mice exhibited progressive motor deficits, and many of these motor abnormalities are due to nigrostriatal dysfunction, we further aimed to explore the probable underlying cause. Because Ube3a has been shown to promote the proteasome-mediated degradation of α-synuclein[29], we intended to investigate the possible implications of this intrinsically disordered protein for nigrostriatal dopaminergic function in AS mice. We first measured α-synuclein levels in the striatum of AS mice and age-matched wild-type animals at postnatal days 25 and 90 (P25 and P90). Surprisingly, we observed a significantly elevated level of α-synuclein in the striatal tissues of AS mice in comparison with wild-type controls at both P25 and P90 (Fig.2A and B). The level of α-synuclein in AS mice was elevated to about 2.5-fold relative to wild-type groups at P25, and further increased to around 3.5-fold at P90. We next analysed the level of the pS129 variant of α-synuclein in these mouse samples. The pS129 derivative of α-synuclein is commonly considered a surrogate marker of pathology in PD and other synucleinopathies, and is also reported to be involved in synaptic transmission [31]. The level of pS129 derivative of α-synuclein was very low and nearly similar in the striatal tissues of both wild-type and AS mice. However, its level was significantly elevated (more than 2-fold) in the striatum of AS mice at P90 compared with the age-matched wild-type group (Fig.2A and C). Since striatal samples of AS mice at P90 displayed considerably increased levels of both normal and pS129 variant α-synuclein, and pS129 variant is associated with insoluble α-synuclein, we further analysed the accumulation of insoluble α-synuclein in these samples. As shown in Figure 2B, the insoluble pool of α-synuclein is significantly increased in the AS mice striatum at P90 (Fig.2D and E). The immunofluorescence localization study further demonstrated increased α-synuclein levels in the substantia nigra dopaminergic neurons of AS mice at P90 (Fig.3A). α-synuclein was localized in both the cytoplasmic and nuclear compartments. The level of the pS129 form of α-synuclein was localized predominantly in the nuclear compartments in the dopaminergic and other cortical neurons in both control and AS mice (Fig.3B). However, the fluorescence intensity of pS129-α-synuclein in AS cases (at P90) was considerably stronger and occasionally showed punctate accumulation in the perinuclear regions (Fig.3B).

**Fig. 2.**
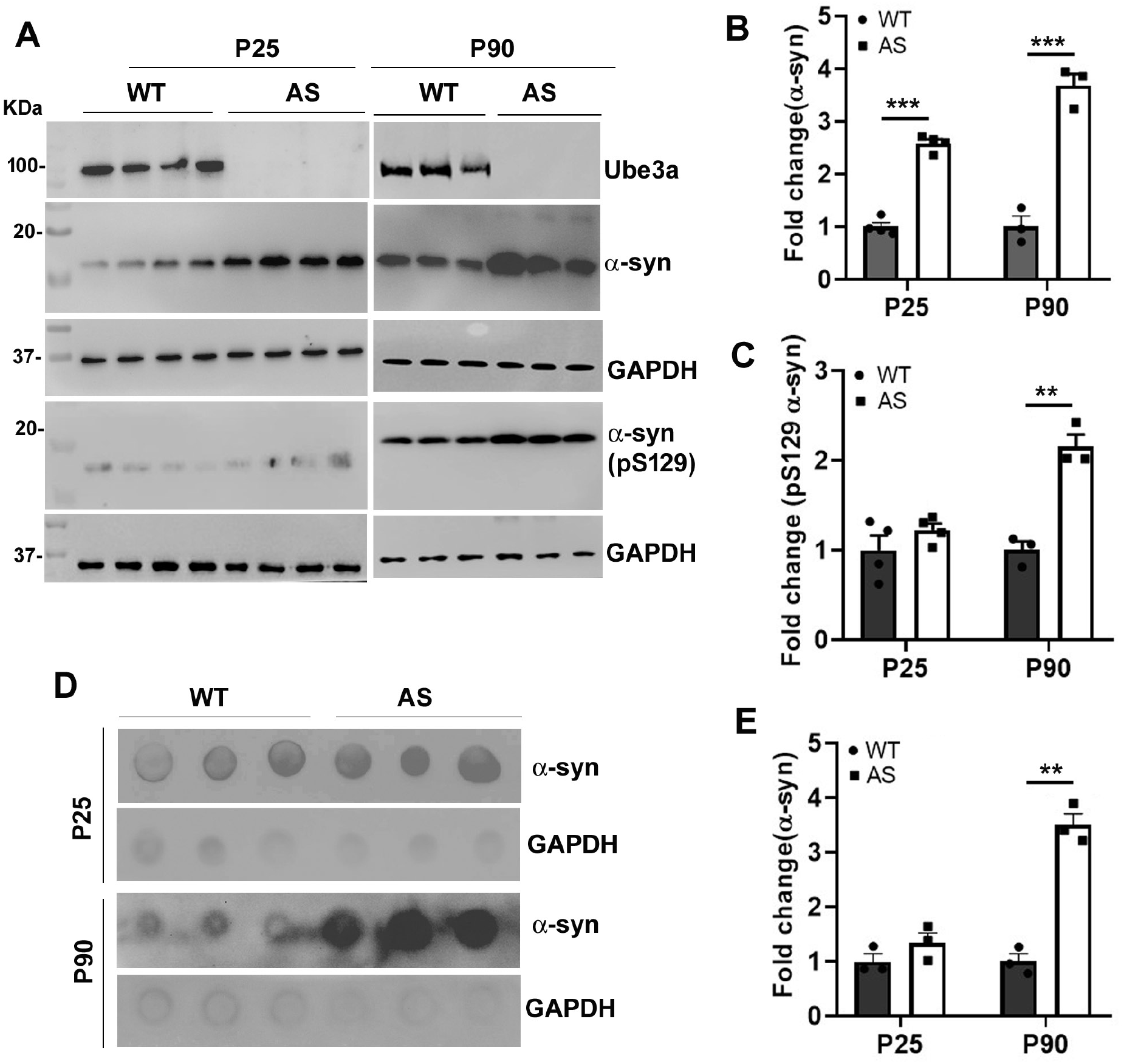
Significant increase in the accumulation of both soluble and insoluble α-synuclein, along with its pS129 variant, in the striatum of AS mice. The striatal tissues from WT and AS animals at P25 and P90 were handled for immunoblot examination utilising antibodies against α-synuclein and its pS129 derivative, Ube3a, and GAPDH. A) Representative immunoblots. B, C) Band intensities of α-synuclein (B) and its pS129 form (C) were computed using NIH Image J software, standardized with GAPDH, and denoted as fold variation. Every track in the blot signifies a sample from a separate mouse. D) The striatal tissues from both WT and AS mice were handled for dot-blot investigation as mentioned in the Materials and Methods and passed through a nitrocellulose membrane in the dot-blot apparatus. Blots were then probed with α-synuclein and GAPDH antibodies. E) The intensities of the dots were measured, standardized with GAPDH, and represented as fold alteration. Data presented are mean ± SEM with samples from three animals in each experimental set at each time point. Values were examined using a mixed-effects analysis, then a post hoc test. \*\*\**P*< 0.001 and \*\**P*< 0.01 related to respective WT group.

**Fig. 3.**
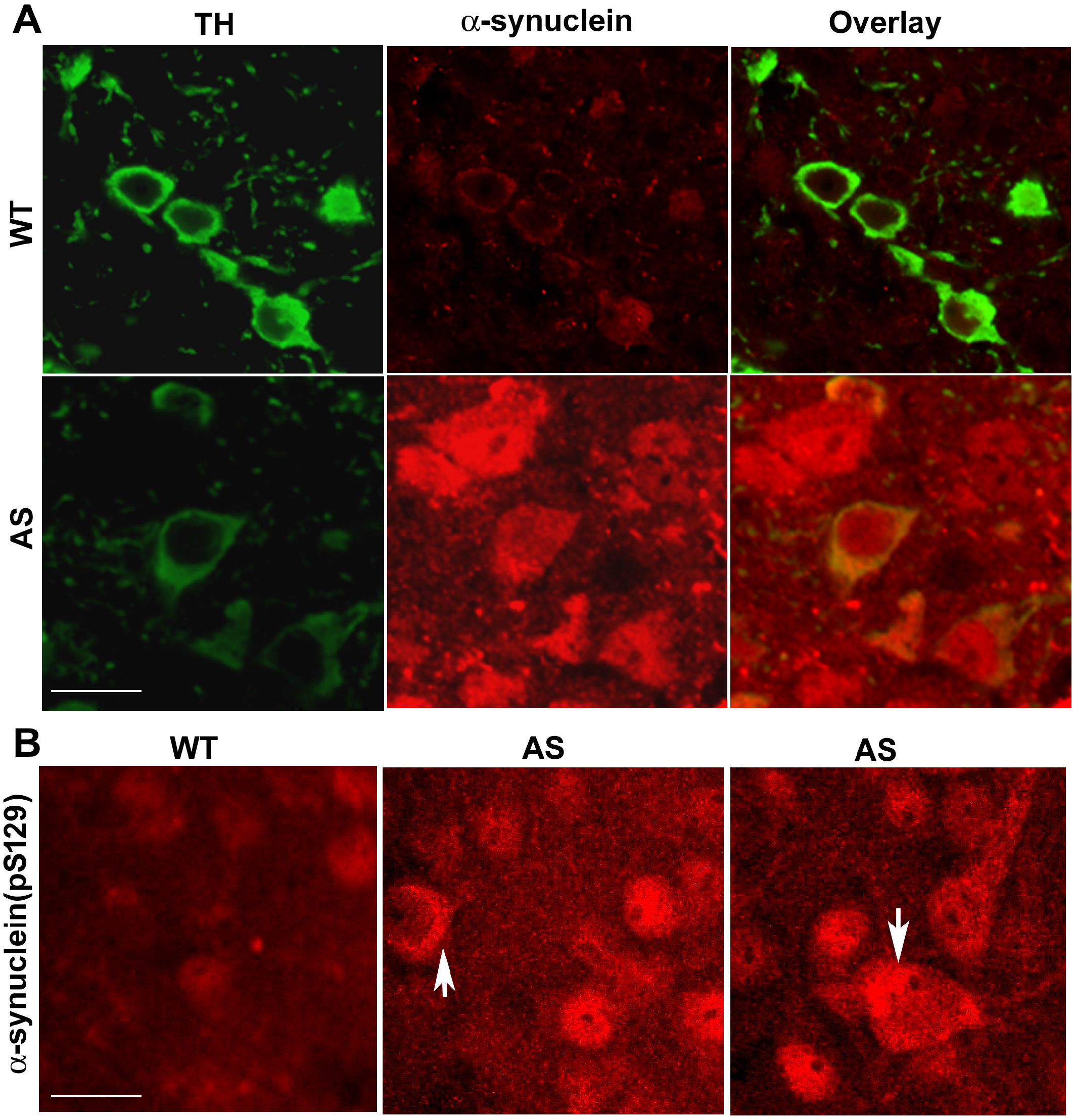
Immunofluorescence localization study shows increased levels of α-synuclein and its pS129 variant in dopaminergic neurons of the substantia nigra and other cortical neurons. A) Coronal brain sections at the midbrain region acquired from both WT and AS mice at P90 were positioned on the same glass slide and continued to a double immunofluorescence study using TH (rabbit specific) and α-synuclein (mouse specific) antibodies. Alexa fluor-594 conjugated secondary antibody was used to visualize α-synuclein, while TH was detected with the use of Alexa fluor-488 tagged secondary antibody. B) Coronal brain sections (midbrain area) from WT and AS mice (P90) were also processed for immunofluorescence staining with pS129 from α-synuclein. Images were taken from the substantia nigra area. Tracing of TH neurons was not possible due to the lack of a rabbit-specific TH antibody. The arrow indicates the accumulation of pS129 derivatives of α-synuclein in the perinuclear region. Three animals from every group were assessed for immunofluorescence staining. Scale bar, 25µm.

### Ube3a interacts with α-synuclein and promotes its K48-linked polyubiquitination

The elevated level of α-synuclein in AS mice brain indicates that Ube3a may target α-synuclein for ubiquitination and subsequent degradation. The K48-linked polyubiquitinated proteins are usually degraded through the proteasome, while the K63-linked polyubiquitinated proteins are targets of autophagic clearance[32,33]. Although Ube3a has been reported to promote proteasomal degradation of α-synuclein in a cell culture-based study, the interaction of Ube3a with α-synuclein and the types of ubiquitin chain formation (K48 or K63) it promotes have not yet been demonstrated. We first studied the interaction between Ube3a and α-synuclein using wild-type and AS mouse cortical lysates in a co-immunoprecipitation assay. Indeed, there is interaction between Ube3a and α-synuclein in the mouse brain lysate (Fig.4A and B). Next, we analysed the K48- and K63-linked polyubiquitination patterns of immunoprecipitated α-synuclein from brain lysates of control and AS mice. The level of K48-linked polyubiquitination was considerably lower in the AS mouse brain lysate than in the wild-type animal (Fig.4D). The K63-linked polyubiquitination of α-synuclein was nearly equal in brain lysates from wild-type and AS mice (Fig.4C). Interestingly, K48-linked total polyubiquitination in AS mouse brain lysate was substantially lower than in wild-type animals, indicating that Ube3a might be targeting proteasome-mediated degradation of many cellular proteins. These results are also in line with our earlier observations that Ube3a promotes proteasome-mediated degradation of α-synuclein.

**Fig. 4.**
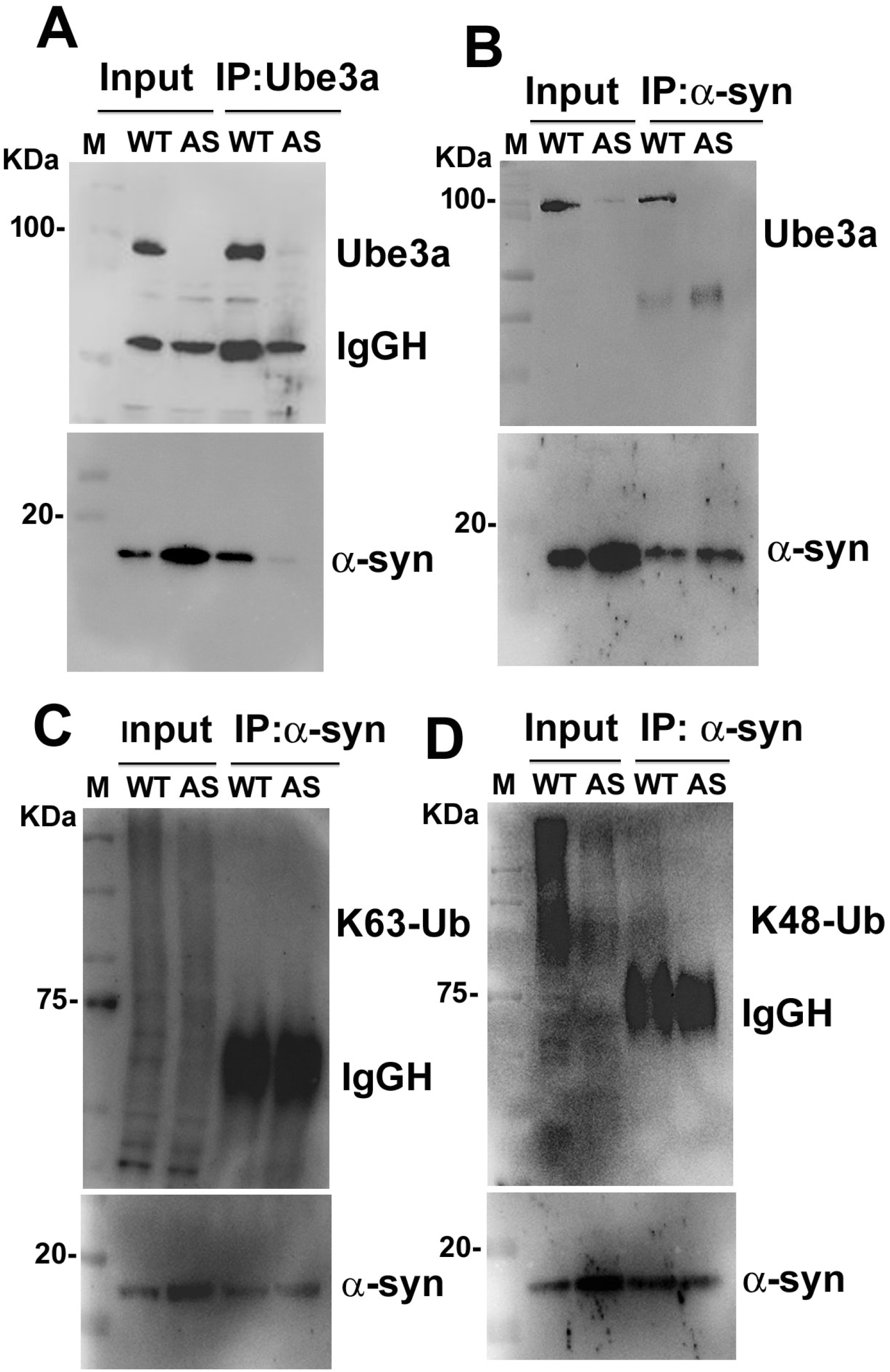
Ube3a interacts with α-synuclein and promotes its K48-linked polyubiquitination. A) Cortical brain lysates from both WT and AS mice were subjected to a co-immunoprecipitation experiment using a Ube3a antibody, and the blots were probed with Ube3a and α-synuclein antibodies. Note the co-immunoprecipitation of α-synuclein by Ube3a in the lysate obtained from WT but not from AS. B) Similar co-immunoprecipitation experiment using α-synuclein antibody, and blots were identified with α-synuclein and Ube3a antibodies. Ube3a was co-immunoprecipitated by α-synuclein. C, D) Cortical lysates were co-immunoprecipitated by α-synuclein antibody, and blots were detected with K63-linked (C) and K48-linked (D) specific polyubiquitin antibodies. K48-linked polyubiquitination of α-synuclein was comparatively lower in the cortical lysate of AS mice. Total K48-linked polyubiquitinated proteins were also considerably lower in the AS mice cortical lysate.

### Altered expression of various dopamine and serotonin pathway-specific genes in the striatum of AS mice at an early developmental period

To investigate the potential impact of the unusually elevated level of α-synuclein in the substantia nigra dopaminergic system of AS mice, we aimed to assess altered expression of dopamine pathway-specific genes in striatal tissues of these animals, compared with age-matched wild-type controls. We used the RT^2^ Profiler PCR Array of mouse dopamine and serotonin pathway-specific genes (84 key genes) involved in the synthesis and metabolism, receptors, transporters, reuptake, and signaling of these neurotransmitters. As shown in Figure 5A, several transcripts showed more than 2-fold alteration in the AS mice striatum in comparison to the control animals at P25. The expression of the up- and down-regulated genes showing about 2-fold alteration was re-verified by RT-qPCR (Fig.5B and C). The Th (Tyrosine hydroxylase, TH), Tph (Tryptophan hydroxylase), Ppp1r1b (Dopamine and cAMP-regulated phosphoprotein 32, Darpp32), Sl6a3 (Dopamine transporter, DAT), Sl18A2 (Vesicular monoamine transporter 2, VMAT2), and Sl6A4 (Serotonin transporter, SERT) transcripts showed significant down-regulation (Fig.5 A). In contrast, Dbh (Dopamine β-hydroxylase), Ptgs2 (Prostaglandin-endoperoxide synthase 2), and Pde4c (Phosphodiesterase 4C) exhibited significant up-regulation in the AS mice striatum at P25 (Fig. 5C). Our findings indicate that the synthesis and transport of both dopamine and serotonin are affected at the adolescent stage in AS mice. The up-regulation of Ptgs2 and Pde4c implied disrupted cAMP-mediated signaling, cellular stress, and inflammation in the brains of AS mice. We further characterized the down-regulated levels of TH and Darpp32 in the striatum of AS mice, along with controls from the P25 and P90 age groups, using immunoblot and immunofluorescence analysis. Both of these proteins showed an age-dependent decrease in their expression in the AS mice striatum in contrast to their respective control group (Fig.6). The Darpp32 showed a dramatic decrease in its expression (in both immunoblot analysis and immunofluorescence study) in AS mice striatum from P25 age, indicating a compromise in dopamine signaling in these animals. This could be due to decreased dopamine synthesis and impaired cAMP signaling. Overall, our results indicate that unusually high levels accumulation of α-synuclein and its pathological variant (pS129) could trigger early dysfunction of the nigrostriatal pathway, resulting in motor deficits, and that the process can be progressive with age (Fig.7).

**Fig. 5.**
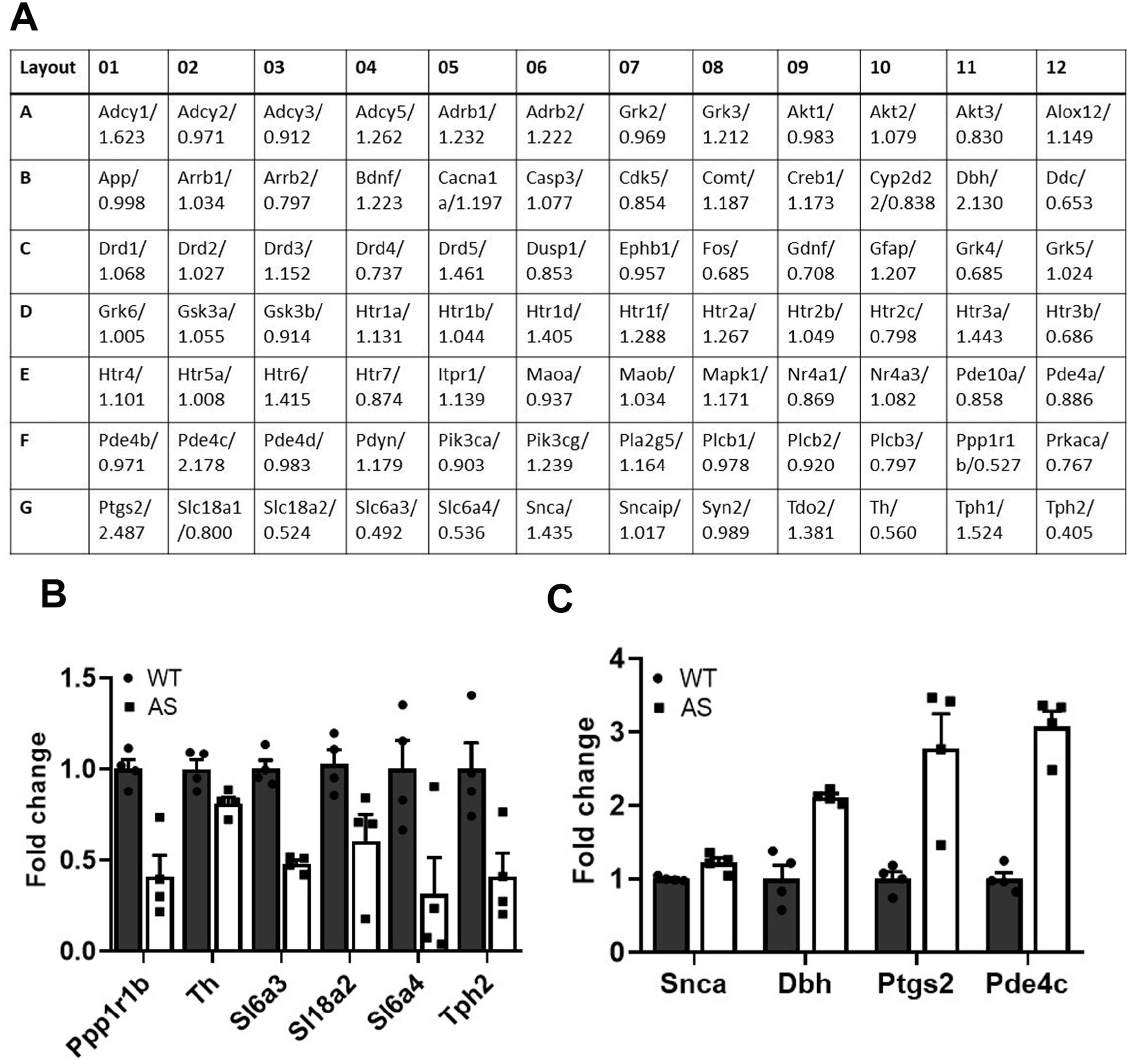
Identification of altered dopamine and serotonin pathway-related genes in the AS mice striatum at P25. Striatal tissues collected from WT and AS animals at P25 were processed for total RNA extraction and then subjected to RT^2^ Profiler PCR Array analysis of dopamine and serotonin path-specific genes as described in the Materials and Methods. A) Typical configuration of an array plate displaying the names of the genes and fold alteration of their transcript levels in AS mice. Striatal tissues from two different animals in each experimental group were used for the array, and average values were used for data analysis. Data were assessed using the GeneGlobe data analysis center (Qiagen) and presented as fold changes. The full names of all the genes, along with references and other experimental quality controls (present in the H row), are shown in Supplementary Table S2. B, C) Verification of the altered gene expression in striatal tissues of AS mice by RT-qPCR. The genes that showed approximately 2-fold alterations in expression on the RT^2^ PCR Profiler array were rechecked. B) Down-regulated genes. C) Up-regulated genes. Values depicted are mean ± SEM with four mouse samples in each set. In B, all transcripts in AS samples are significantly decreased compared to the WT group. In C, except for Snca, all transcripts show up-regulated expression (*t*-test). The *P* values for different transcripts are as follows: Ppp1r1b, *P*=0.003; Th, *P*=0.02; Sl6a3, *P*=0.0011; Sl18a2, *P*=0.036; Sl6a4, *P*=0.035; Tph2, *P*=0.021; Dbh, *P*=0.001; Ptgs2, *P*=0.009; Pde4c, *P*=0.001. Th, Tyrosine hydroxylase; Tph, Tryptophan hydroxylase; Ppp1r1b, Dopamine and cAMP-regulated phosphoprotein 32; Sl6a3, Dopamine transporter; Sl18A2, Vesicular monoamine transporter 2; and Sl6A4, Serotonin transporter; Dbh, Dopamine β-hydroxylase; Ptgs2, Prostaglandin-endoperoxide synthase 2; and Pde4c, Phosphodiesterase 4C; Snca, α-synuclein.

**Fig. 6.**
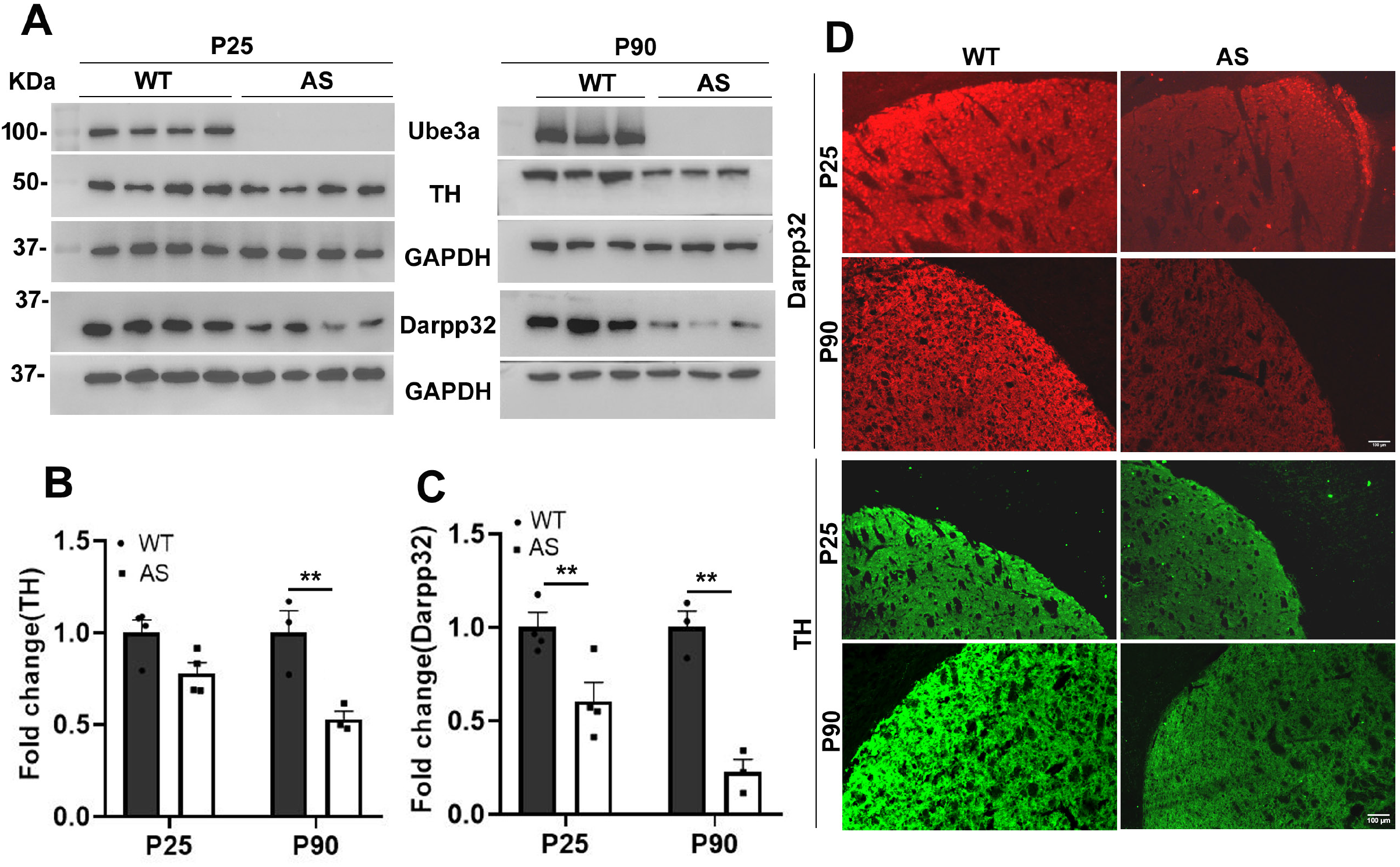
Age-related down-regulation of TH and Darpp32 in the striatum of AS mice. **A, B, C)** Striatal tissues obtained from WT and AS mice at P25 and P90 were handled for immunoblot study utilising antibodies against TH, Darpp32, Ube3a, and GAPDH. Band densities of the blots were measured, standardized against GAPDH, and shown as fold alteration. A) Immunoblots. A sample from a different mouse was used in each lane. B, C) Age-dependent down-regulation of TH and Darpp32. Values shown are mean ± SEM with tissues from three different mice in each set. The data were assessed applying mixed-effects analysis, preceded by a post hoc test. D) Immunofluorescence staining of TH and Darpp32 showing their down-regulation in the striatum of AS mice in comparison with the WT set. \*\**P*< 0.01 and \**P*< 0.05 related to respective WT set. Brain sections from the midbrain area (including the striatal region) of different experimental group were mounted on the same slide and processed for immunofluorescence procedure with TH and Darpp32 primary antibodies, preceded by Alexa Fluor-488-tagged secondary antibody. Typical images of TH and Darpp32 are presented. Three animals from each experimental set were analysed for the immunofluorescence study. Scale bar: 100 µm. Arrow indicates the punctate staining.

**Fig. 7.**
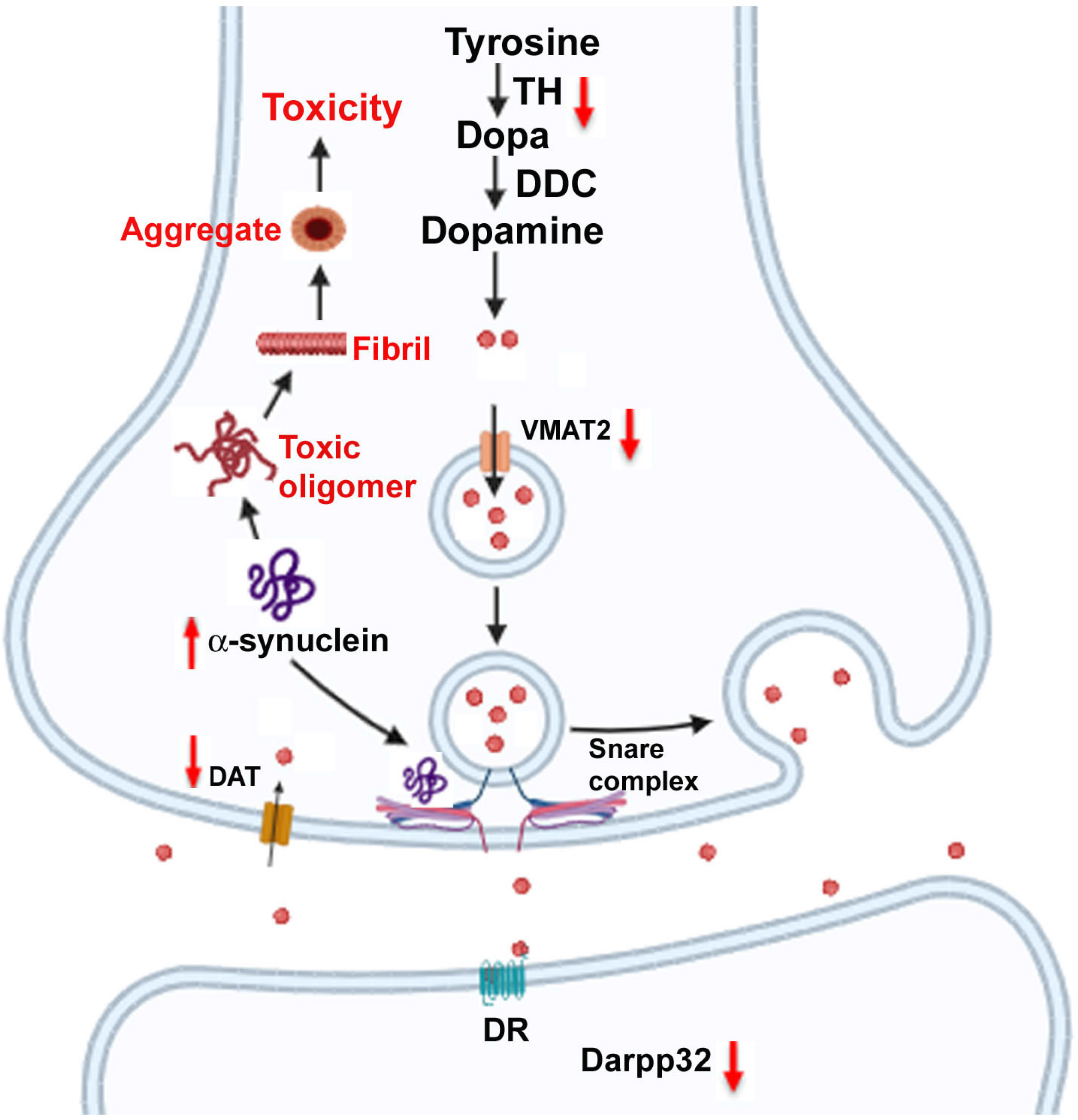
Proposed mechanisms of α-synuclein-induced nigrostriatal dysfunction in AS mice. Under normal circumstances, α-synuclein helps SNARE complex assembly, assists in synaptic vesicle docking, and dopamine release. However, lack of Ube3a in AS mice causes increased accumulation of α-synuclein, leading to its misfolding and the development of toxic oligomers, fibrils, and aggregates. The aberrant accumulations of various abnormal forms could eventually disrupt synaptic transmission and neuronal function. The red-marked arrow and text indicate changes in the AS condition. Picture is drawn with the help of BioRender.

## Discussion

Accumulations of α-synuclein into insoluble aggregates are the classic pathological hallmark of synucleinopathies, a group of neurodegenerative disorders comprising PD, dementia with Lewy bodies, and multiple system atrophy[34–37]. α-synuclein is a naturally disordered protein and is normally involved in synaptic vesicle trafficking and neurotransmitter release[38]. However, duplication, triplication, or gain-of-function mutations of the *SNCA* gene accelerate the production, misfolding, and aggregation of α-synuclein, which eventually initiate neuronal dysfunction and neurodegeneration[39,35]. Here, we report for the first time that loss of Ube3a function in AS mice results in an aberrant accumulation of α-synuclein in their brains and that this accumulation may be linked to the pathogenesis of AS.

Although an earlier study reported degradation of α-synuclein by Ube3a, the interaction between Ube3a and α-synuclein, the degradation mechanisms, and, most importantly, the implication of α-synuclein in the pathogenesis of AS had not been investigated. We first observed age-dependent increases in soluble and insoluble α-synuclein and its pathological variant (pS129) in the striatum of AS mice. Our results also showed that Ube3a interacts with α-synuclein and promotes its K48-linked polyubiquitination. The Ube3a-deficient mouse brain exhibits reduced levels of total K48-linked polyubiquitination and K48-linked polyubiquitinated α-synuclein. These results advocate that Ube3a act as a crucial player in regulating the proteasome-mediated degradation of α-synuclein, since proteins modified by K48-linked polyubiquitination are primarily targeted for clearance by the proteasome [32]. Interestingly, significantly reduced levels of total and K48-linked ubiquitination in the brains of AS mice suggest that Ube3a may target the proteasomal degradation of many other substrates that are yet to be identified. Several ubiquitin ligases have been found to promote the degradation of α-synuclein through either proteasome or autophagy, including Nedd4, SIAH, SCF-FBXL5, Parkin, CHIP, and listerin[40–43]. However, Ube3a seems unique for several reasons. First of all, α-synuclein pathology and its link to disease progression, particularly progressive motor deficits, in AS mice are novel. Secondly, Ube3a could be a crucial modifier of synucleinopathies. Thirdly, Reactivation of paternal Ube3a in the neuron might provide an innovative therapeutic approach for synucleinopathies and other neurodegenerative disorders characterized by significantly reduced Ube3a levels[44,45]. The reactivation of paternal Ube3a through small molecules or Ube3a-ATS antisense oligonucleotides is considered one of the most promising strategies for treating AS[46,47]. Similar strategies could be useful in synucleinopathies and other neurodegenerative disorders.

Another interesting observation was the down-regulation of selected dopamine and serotonin pathway-specific genes, such as TH, Darpp32, DAT, VMAT2, SERT, and Tph2 in the striatum of AS mice at a very early age, i.e., at P25. All these dopamine and serotonin pathway-specific genes were known to be decreased in post-mortem brain tissues from PD and also in the striatal regions of various animal models of PD[48,49,35,50,51]. In various α-synuclein models of PD, pathological accumulation or overexpression of misfolded α-synuclein triggers a cascade of events, including mitochondrial dysfunction, oxidative stress, and epigenetic alterations, that eventually lead to decreased expression of all these dopamine pathway-specific genes and induce dysfunction of dopaminergic neurons[35,48,50]. Therefore, it is conceivable that increased accumulation of α-synuclein in the substantia nigra dopaminergic neurons/striatum of AS mice could be directly linked with the down-regulation of all these dopamine and serotonin pathways-specific genes, and altered expression of all these genes could be the underlying cause of progressive nigrostriatal dysfunction and motor deficits observed in these animals. Mitochondrial dysfunction and elevated reactive oxygen species were detected in the hippocampus of AS mice and in AS brain-derived neuronal precursor cells, which may be due to increased α-synuclein accumulation[52,53]. Furthermore, up-regulation of Ptgs2 in the striatum of AS mice indicates that neurons are under stress, which in turn might stimulate an inflammatory response. The augmented expression of Pde4c in the striatum of AS mice suggests disrupted cAMP-mediated signaling and provides a rationale for Darpp32 down-regulation. The down-regulation of Tph and SERT indicates defective serotonin synthesis and reuptake in the AS mice brain and could explain why some selective serotonin reuptake inhibitors reduce hyperactivity and anxiety in some AS patients and also in a mouse model of AS[54,55]. How TH and Tph are down-regulated in AS mouse brains is unclear, but might be linked to decreased expression of GTP cyclohydrolase-1, a rate-limiting enzyme in monoamine synthesis[28].

AS patients (a 23-year-old male and a 43-year-old female) have been revealed to exhibit distinctive features of PD, such as tremors, cogwheel rigidity, and bradykinesia, and these motor deficits responded well to levodopa[24]. Further studies demonstrated that the motor deficits exhibited by relatively older AS mice are due to dysfunctional nigrostriatal dopaminergic system[25,26]. Surprisingly, a double-blind randomized clinical trial in children with AS (4-12 years old) treated with levodopa failed to improve motor deficits[27]. The reason for the failure of the levodopa clinical trial is unclear and must be critically analysed. Although, the dose of levodopa and the sensitivity of the end-point measures were raised as probable factors, the age of the recruited AS patients could also influence the outcome. Significant nigrostriatal dysfunction in AS patients might take place at comparatively older ages.

Until now, numerous substrates of Ube3a have been identified, and many of them are directly or indirectly linked in modulating synaptic activity and plasticity and connected in AS pathogenesis[21,20,56,57,16,19]. α-synuclein, under normal conditions, regulates synaptic vesicle trafficking to the pre-synaptic membrane and fine-tunes neurotransmitter release[38]. However, its high local concentration in neurons drives structural destabilization, followed by misfolding and the formation of toxic oligomers that could eventually disrupt synaptic function and communication, leading to cognitive, motor, and psychiatric symptoms frequently observed in synucleinopathies [35,39]. Triplications of *SNCA* gene are reported to cause early onset PD with dementia, while its duplications lead late onset PD that progresses slowly without dementia [58]. Our results indicate that some of the typical features of synucleinopathies are also present in AS mice. Our study focused on the impact of abnormally accumulated α-synuclein on nigrostriatal pathway dysfunction and related motor deficits in AS mice. It would also be interesting to see the influence of abnormally accumulated α-synuclein on the intellectual and other behavioral discrepancies in these animals. Our results also prompted us to speculate that Ube3a dysfunction could be linked to proteostasis failure and the progression of PD and related synucleinopathies. Currently, there is no reported genetic association between *UBE3A* and synucleinopathies. However, oxidative stress appears to alter the solubility and activity of Ube3a across various cellular and animal models of neurodegenerative diseases[59,45,44]. Therefore, an age-dependent decrease in Ube3a function might contribute to the progressive phenotypic features of synucleinopathies.

Overall, our findings highlight that Ube3a is a novel regulator of α-synuclein, and abnormal accumulations of α-synuclein in the brains of AS mice could be linked to the progressive motor and other behavioral insufficiencies observed in these animals. Our results also speculate that Ube3a may be a new modifier of synucleinopathies.

## Acknowledgements

We would like to sincerely thank all the laboratory members for the discussion and input.

## Compliance with ethical standards

All animal investigations were carried out in agreement with the stringent rules defined by the Committee for the Purpose of Control and Supervision of Experiments on Animals, Ministry of Environment and Forestry, Government of India and were permitted by the Institutional Animal Ethics Committee of the Indian Institute of Technology Kharagpur (Protocol number: IE-3/NJ-BS/1.21).

## Data availability statement

Data will be made available by the corresponding author on request.

## Author contributions

All authors contributed study conception and design. Experimentations were performed by Sagarika Das and Bhaskarjyoti Giri; data were analysed by Sagarika Das and Nihar Ranjan Jana. First draft of the manuscript was written by Nihar Ranjan Jana and all authors read, commented and approved the final manuscript.

