## Supplementary information for "Aberrant accumulation of α-synuclein might be linked with the progressive motor deficits in a mouse model of Angelman syndrome"

**Supplementary Table 1**

**List of primers used in the experiment**

| Gene | | Primer Sequence(5'-3') |
|---|---|---|
| Dbh | FP | GGCTCTGTATGACTACGCCC |
| | RP | GGCTTCTCCGTTCAGTTGGA |
| Slc6a3 | FP | CAAAGCTGAAGTCTGACGCTGG |
| | RP | AGAAGACATTGGTCCCACGGAG |
| Tph2 | FP | CTGAATCCGCCTGAGAGCAT |
| | RP | CCGTACATGAGGACTCGGTG |
| Snca | FP | CAGCAGTCGCTCAGAAGACA |
| | RP | AACTGAGCACTTGTACGCCA |
| Slc6a4 | FP | GCCATCAGCCCTCTGTTTCT |
| | RP | TGGCTTAGAGGGGAGGAGTC |
| Th | FP | GGGCTGCTGTCTTCCTATGG |
| | RP | TACACCGGCTGGTAGGTTTG |
| Ptgs2 | FP | AGCCCATTGAACCTGGACTG |
| | RP | ACCCAATCAGCGTTTCTCGT |
| Pde4c | FP | CGGAACCTCAGTACCAAGCA |
| | RP | GCCAAGGCTGGTCACTTTCT |
| Sl18a2 | FP | AACACGACTGTCCCTCCCGACT |
| | RP | GGCGATCAGCAGGAAGGCATAG |
| Ppp1r1b | FP | CCCAAAGTCGAAGAGACCCA |
| | RP | CCGAAGCTCCCCTAACTCATC |
| GAPDH | FP | ATGACTCTACCCACGGCAAG |
| | RP | CTGGAAGATGGTGATGGGTT |

**Supplementary Table2**

| Well Posi | Gene Name | Symbol | Fold Change |
|---|---|---|---|
| A01 | Adenylate cyclase 1 | Adcy1 | 1.623932157 |
| A02 | Adenylate cyclase 2 | Adcy2 | 0.971374047 |
| A03 | Adenylate cyclase 3 | Adcy3 | 0.912037061 |
| A04 | Adenylate cyclase 5 | Adcy5 | 1.262694909 |
| A05 | Adrenergic receptor, beta 1 | Adrb1 | 1.232124603 |
| A06 | Adrenergic receptor, beta 2 | Adrb2 | 1.222265689 |
| A07 | Adrenergic receptor kinase, beta 1 | Adrbk1 | 0.969589955 |
| A08 | Adrenergic receptor kinase, beta 2 | Adrbk2 | 1.21225793 |
| A09 | Thymoma viral proto-oncogene 1 | Akt1 | 0.98269872 |
| A10 | Thymoma viral proto-oncogene 2 | Akt2 | 1.079852037 |
| A11 | Thymoma viral proto-oncogene 3 | Akt3 | 0.829777102 |
| A12 | Arachidonate 12-lipoxygenase | Alox12 | 1.148635192 |
| B01 | Amyloid beta (A4) precursor protein | App | 0.99821678 |
| B02 | Arrestin, beta 1 | Arrb1 | 1.033606556 |
| B03 | Arrestin, beta 2 | Arrb2 | 0.797340821 |
| B04 | Brain derived neurotrophic factor | Bdnf | 1.222905137 |
| B05 | Calcium channel, voltage-dependent, P/Q type, alpha | Cacna1a | 1.197364438 |
| B06 | Caspase 3 | Casp3 | 1.07717289 |
| B07 | Cyclin-dependent kinase 5 | Cdk5 | 0.854138822 |
| B08 | Catechol-O-methyltransferase | Comt | 1.187113479 |
| B09 | CAMP responsive element binding protein 1 | Creb1 | 1.173039242 |
| B10 | Cytochrome P450, family 2, subfamily d, polypeptide 2 | Cyp2d22 | 0.838164315 |
| B11 | Dopamine beta hydroxylase | Dbh | 2.130237698 |
| B12 | Dopa decarboxylase | Ddc | 0.653314438 |
| C01 | Dopamine receptor D1A | Drd1a | 1.068492161 |
| C02 | Dopamine receptor D2 | Drd2 | 1.02680448 |
| C03 | Dopamine receptor D3 | Drd3 | 1.152834461 |
| C04 | Dopamine receptor D4 | Drd4 | 0.737158903 |
| C05 | Dopamine receptor D5 | Drd5 | 1.461209184 |
| C06 | Dual specificity phosphatase 1 | Dusp1 | 0.853759787 |
| C07 | Eph receptor B1 | Ephb1 | 0.957989616 |
| C08 | FBJ osteosarcoma oncogene | Fos | 0.685407767 |
| C09 | Glial cell line derived neurotrophic factor | Gdnf | 0.708042851 |
| C10 | Glial fibrillary acidic protein | Gfap | 1.207038466 |
| C11 | G protein-coupled receptor kinase 4 | Grk4 | 0.685165538 |
| C12 | G protein-coupled receptor kinase 5 | Grk5 | 1.024472407 |
| D01 | G protein-coupled receptor kinase 6 | Grk6 | 1.005709211 |
| D02 | Glycogen synthase kinase 3 alpha | Gsk3a | 1.055438091 |
| D03 | Glycogen synthase kinase 3 beta | Gsk3b | 0.914057921 |
| D04 | 5-hydroxytryptamine (serotonin) receptor 1A | Htr1a | 1.13130393 |
| D05 | 5-hydroxytryptamine (serotonin) receptor 1B | Htr1b | 1.044360401 |
| D06 | 5-hydroxytryptamine (serotonin) receptor 1D | Htr1d | 1.405525482 |
| D07 | 5-hydroxytryptamine (serotonin) receptor 1F | Htr1f | 1.288217232 |
| D08 | 5-hydroxytryptamine (serotonin) receptor 2A | Htr2a | 1.267951948 |
| D09 | 5-hydroxytryptamine (serotonin) receptor 2B | Htr2b | 1.049506802 |
| D10 | 5-hydroxytryptamine (serotonin) receptor 2C | Htr2c | 0.798451651 |
| D11 | 5-hydroxytryptamine (serotonin) receptor 3A | Htr3a | 1.443971955 |
| D12 | 5-hydroxytryptamine (serotonin) receptor 3B | Htr3b | 0.686376134 |
| E01 | 5 hydroxytryptamine (serotonin) receptor 4 | Htr4 | 1.101732447 |
| E02 | 5-hydroxytryptamine (serotonin) receptor 5A | Htr5a | 1.008598348 |
| E03 | 5-hydroxytryptamine (serotonin) receptor 6 | Htr6 | 1.415243711 |

| | | | |
|---|---|---|---|
| E04 | 5-hydroxytryptamine (serotonin) receptor 7 | Htr7 | 0.874992469 |
| E05 | Inositol 1,4,5-trisphosphate receptor 1 | Itpr1 | 1.139188607 |
| E06 | Monoamine oxidase A | Maoa | 0.937319086 |
| E07 | Monoamine oxidase B | Maob | 1.034467539 |
| E08 | Mitogen-activated protein kinase 1 | Mapk1 | 1.171339285 |
| E09 | Nuclear receptor subfamily 4, group A, member 1 | Nr4a1 | 0.869869591 |
| E10 | Nuclear receptor subfamily 4, group A, member 3 | Nr4a3 | 1.082616802 |
| E11 | Phosphodiesterase 10A | Pde10a | 0.858823882 |
| E12 | Phosphodiesterase 4A, cAMP specific | Pde4a | 0.886554791 |
| F01 | Phosphodiesterase 4B, cAMP specific | Pde4b | 0.971325862 |
| F02 | Phosphodiesterase 4C, cAMP specific | Pde4c | 2.178370653 |
| F03 | Phosphodiesterase 4D, cAMP specific | Pde4d | 0.983036281 |
| F04 | Prodynorphin | Pdyn | 1.179640193 |
| F05 | Phosphatidylinositol 3-kinase, catalytic, alpha polypept | Pik3ca | 0.903020734 |
| F06 | Phosphoinositide-3-kinase, catalytic, gamma polypepti | Pik3cg | 1.239211919 |
| F07 | Phospholipase A2, group V | Pla2g5 | 1.164853307 |
| F08 | Phospholipase C, beta 1 | Plcb1 | 0.978829058 |
| F09 | Phospholipase C, beta 2 | Plcb2 | 0.920615145 |
| F10 | Phospholipase C, beta 3 | Plcb3 | 0.797207206 |
| F11 | Protein phosphatase 1, regulatory (inhibitor) subunit 1I | Ppp1r1b | 0.527300213 |
| F12 | Protein kinase, cAMP dependent, catalytic, alpha | Prkaca | 0.767946911 |
| G01 | Prostaglandin-endoperoxide synthase 2 | Ptgs2 | 2.487589458 |
| G02 | Solute carrier family 18 (vesicular monoamine), memb | Slc18a1 | 0.800778279 |
| G03 | Solute carrier family 18 (vesicular monoamine), memb | Slc18a2 | 0.524017726 |
| G04 | Solute carrier family 6 (neurotransmitter transporter, do | Slc6a3 | 0.492935726 |
| G05 | Solute carrier family 6 (neurotransmitter transporter, se | Slc6a4 | 0.536122516 |
| G06 | Synuclein, alpha | Snca | 1.435032343 |
| G07 | Synuclein, alpha interacting protein (synphilin) | Sncaip | 1.01757434 |
| G08 | Synapsin II | Syn2 | 0.989909237 |
| G09 | Tryptophan 2,3-dioxygenase | Tdo2 | 1.381414491 |
| G10 | Tyrosine hydroxylase | Th | 0.560064439 |
| G11 | Tryptophan hydroxylase 1 | Tph1 | 1.524915337 |
| G12 | Tryptophan hydroxylase 2 | Tph2 | 0.405956213 |
| H01 | Actin, beta | Actb | 1.086474411 |
| H02 | Beta-2 microglobulin | B2m | 0.764133196 |
| H03 | Glyceraldehyde-3-phosphate dehydrogenase | Gapdh | 1 |
| H04 | Glucuronidase, beta | Gusb | 0.844629247 |
| H05 | Heat shock protein 90 alpha (cytosolic), class B memb | Hsp90ab1 | 0.965622794 |
| H06 | Mouse Genomic DNA Contamination | MGDC | #VALUE! |
| H07 | Reverse Transcription Control | RTC | 1.277466549 |
| H08 | Reverse Transcription Control | RTC | 1.13923138 |
| H09 | Reverse Transcription Control | RTC | 0.962574828 |
| H10 | Positive PCR Control | PPC | 0.98302708 |
| H11 | Positive PCR Control | PPC | 0.850106561 |
| H12 | Positive PCR Control | PPC | 0.94144255 |
